# Electrophysiological correlates of cognitive reserve in healthy older adults at different cognitive states

**DOI:** 10.1101/2024.12.30.627948

**Authors:** Alina Zhunussova, Maria Concetta Pellicciari, Manuel Carreiras, Jesús Cespón

**Author notes:** **Corresponding author**: Alina Zhunussova Email address, Postal address: Universitätsstrasse 5-7, 6020, Innsbruck, Austria, -: 56092.

## Abstract

High cognitive reserve (CR) is associated with a set of lifestyle factors such as high education level and occupational attainment. Studies suggest that CR is linked to enhanced executive control functions. Nevertheless, it is unclear what are the functional neural activity patterns linked to high CR and to what extent they are steady across different cognitive states. The aim of the present study was to investigate electrophysiological differences between low CR (LCR) and high CR (HCR) at rest and during the performance of two cognitive control tasks with different difficulty level. Thus, 67 older adults performed an experimental session consisting of two spatial stimulus-response compatibility (SRC) tasks (i.e., the Simon task, and the more demanding spatial Stroop task), and a resting state during an electroencephalogram (EEG) recording. Cognitive control was better in HCR than LCR group, as revealed by higher accuracy in HCR compared to LCR group in incongruent conditions of the tasks. Moreover, event-related brain potentials (ERP) showed earlier latencies (P300) in HCR than LCR and time/frequency analysis showed higher alpha activity in HCR than LCR during the performance of the tasks, while no differences were observed at rest within any of the analyzed frequency bands. Higher P300 frontalization was observed in LCR than HCR in the more difficult task but it was not related to frontalization within a specific frequency band. Overall, results showed that electrophysiological differences between high and low CR depend on the cognitive state, with greater CR-related differences emerging at increased task demands.

## 1. INTRODUCTION

Considering the increased aging of the population all over the world, studying variables and mechanisms related to successful cognitive aging is an important research topic. The construct of cognitive reserve (CR) refers to the brain capability to implement compensatory mechanisms in response to brain damage underlying the physiological aging or a pathological condition (Stern, 2009). For instance, for the same degree of neural damage related to Alzheimer’s disease (AD), older adults with high CR exhibit less clinical symptoms compared to older adults with low CR (Darby et al., 2017). Thus, investigating neural activity patterns associated with high CR is a relevant issue because it may reveal the neural activity patterns that should be induced after applying interventional programs aimed to improve cognition and delay the onset of dementia.

CR is the result of a complex set of biological and environmental factors that are associated with the optimization of brain functioning in older adults (Stern, 2009). There is not a direct way to measure CR, but it can be estimated by means of proxy variables (e.g., years of education, occupational status) and several lifestyles and cognitively stimulating activities such as physical exercise, playing musical instruments or speaking more than one language (Tucker & Stern, 2011; Stern et al., 2020). According to longitudinal studies, individuals with high educational level and occupational attainment are at a lower risk of developing dementia (Dekhtyar et al., 2016; Almeida-Meza et al., 2021). Research in older adults has suggested that executive functions, which allow carrying out goal directed actions (Miller & Cohen, 2001; Miyake et al., 2000), are enhanced in high compared to low CR individuals (Cabral et al., 2016; Darby et al., 2017; Roldán-Tapia et al., 2012). These results are highly relevant considering that executive functions substantially decline with aging (Grady, 2012; Ferguson et al., 2021; Zacks, 2000) as a result of physiological changes within frontal and parietal areas (Bagarinao et al., 2022).

The stimulus-response compatibility (SRC) paradigm has been widely used to study executive functions. During the performance of SRC tasks, participants have to suppress the bias to respond on the basis of a salient but irrelevant stimulus feature in order to implement an alternative response based on a relevant stimulus feature, which is previously defined in the task instructions. Two typical examples of SRC tasks are the spatial Stroop task and the Simon task. Several studies support the notion that Simon and Stroop tasks represent one-model tasks that share an underlying mechanism for resolving cognitive interference, as both require the suppression of irrelevant information in order to respond to task-relevant features (Faber et al., 1986; Hasbroucq & Guiard., 1991; Kornblum et al., 1999; Zhang et al., 1999). Whereas in the Simon task there is an incongruence between the stimulus location and response side (i.e., stimulus-response (S-R) conflict), in the spatial Stroop task the incongruence is between stimulus location and response side and additionally between the direction pointed by the arrow and its location (i.e., S-R and S-S conflicts) (Cespón et al., 2020). These two sources of conflict make the spatial Stroop task a more demanding task compared to the Simon task (Cespón et al., 2023; Juncos-Rabadán et al., 2008). An early study by O’Leary and Barber (1993) had confirmed that the spatial Stroop task causes more significant spatial interference and results in poorer performance in comparison to the Simon task. Considering that difficulty of the task has been suggested as an experimental factor linked to the appearance of differences between low and high CR groups (Martinez et al., 2022), having a sample of participants performing two similar tasks with different difficulty levels (i.e., spatial Stroop and Simon tasks), is an appropriate approach to test to what extent neural activity differences between high and low CR depend on the brain state or whether the difficulty of the task represents a key variable to observe such differences.

The high temporal resolution of electroencephalogram (EEG) and event-related potentials (ERP) is useful to investigate neural correlates of cognitive processes that could not be isolated by using behavioral measures. Previous studies suggest that ERP analyses could be useful to isolate specific cognitive control differences between high and low CR during the performance of SRC tasks (Cespón et al., 2020; Cespón & Carreiras, 2020). Also, analyses of the EEG resting state help us to investigate, in a more comprehensive way, to what extent differences between high and low CR depend on the cognitive state.

The N200 and P300 components are commonly employed ERPs for investigating the electrophysiological correlates of executive functions (Cespón et al., 2020; Cespón et al., 2023). The N200 is a negative ERP component that appears approximately between 200 ms and 350 ms and the N200 waveform related to cognitive control is primarily distributed in the anterior brain regions (i.e., fronto-central N200) (Folstein & Van Petten, 2008). The N200 modulations have been associated with inhibition, conflict, and error monitoring (Folstein & Van Petten, 2008), particularly increasing in amplitude for trials with conflicting information compared to trials with no conflicting information during the performance of SRC tasks (Cespón & Carreiras, 2020; Heil et al., 2000; Kopp et al., 1996; Larson et al., 2014). The P300 is a positive ERP component, peaking between 300 and 600 ms after stimulus onset (Polich, 2007). During the performance of SRC tasks, P300 is a correlate of stimulus-response binding update and its latency its usually later in incongruent (or conflicting) compared to congruent (or no conflicting) trials (Leuthold & Sommer, 1999; Leuthold & Schröter, 2006; Melara et al., 2008). In addition, P300 ERP is usually lower in conflicting than non-conflicting trials (Cespón et al., 2013; Leuthold, 2011). However, not only do conflict conditions influence these ERP correlates, but also physiological aging significantly modulates them, generally resulting in extended latencies and reduced amplitudes (Amenedo et al., 2012; Cespón & Carreiras, 2020; Cespón et al., 2013; Gajewski et al., 2018). As an example, in interference tasks, Clawson et al. (2017) reported a diminished distinction in amplitudes between conflicting and non-conflicting conditions. Another recent study has shown that individuals with high СR exhibit neural activity patterns similar to those observed in young adults, characterized by earlier latencies and heightened amplitudes (Cespón et al., 2023).

In parallel with event-related potential studies, a growing body of research has focused on rhythmic oscillatory activity. Multiple studies have investigated how rhythmic oscillatory activity during resting state and tasks performance relates to cognitive functioning in healthy adults. Previous studies suggested that increased frontal theta activity is connected to better working memory (Itthipuripat et al., 2013; Klimesch et al., 2005; Pavlov & Kotchoubey, 2017; Zakrzewska & Brzezicka, 2014). Likewise, studies have shown that frontal midline theta power increases during higher cognitive workload (Gevins & Smith, 2000) in tasks involving working memory (Mitchell et al., 2008) and executive functions (Nigbur et al., 2011). In this context, alpha activity was related to reactivation of memory codes in working memory (Klimesch et al., 2005). Beta activity was related to underline high-level mental processes, particularly in sustaining the cognitive mindset and prioritizing top-down influences over new or surprising external stimuli (Engel & Fries, 2010). Aging in general was related to reduced theta and alpha activity in addition to increased activity in the beta frequency band (Barry & De Blasio, 2017). Nevertheless, there is not a clear understanding about the relationships between brain oscillations and CR in older adults (Balart-Sánchez et al., 2021).

Studies in the field of neurocognitive aging have demonstrated that increased age is accompanied by changed brain activity patterns during the cognitive performance, mainly characterized by a posterior to anterior shift of activity and inter-hemispheric differentiation (McDonough et al., 2022). A recent ERP study analyzing the P300 component suggested that high CR is associated with maintenance of neural activity patterns observed in younger adults, preventing or minimizing the posterior-to-anterior shift of brain activity manifested by lower CR older adults during the performance of cognitive control tasks (Cespón et al., 2023). A pending research question is whether there is a specific frequency band contributing to posterior to anterior shift of activity during the performance of cognitive control tasks. Studying this issue may allow obtaining more specific electrophysiological correlates of these mentioned age-related changes. In addition, analyzing the EEG spectral activity is a useful approach to explore whether a posterior to anterior shift of activity observed during the cognitive task performance may also occur during the resting state.

The main objective of the current study is to identify neural activity patterns associated with high CR and assess to what extent those correlates are modulated by the cognitive state. Thus, samples of older adults with high and low CR performed two SRC tasks with different difficulty level (i.e., the previously mentioned Simon task and the spatial Stroop task) and a resting state during an EEG recording. EEG spectral power in theta, alpha and beta bands were analyzed at rest and event-related spectral power (ERSP) was calculated during the performance of the tasks. Moreover, during the performance of the mentioned tasks, we have analyzed the N200 and P300 ERP components.

In the present research, we have made the following predictions: 1) we hypothesize that, mainly during the more difficult task (i.e., the spatial Stroop task), ERP latencies will be earlier and larger in high compared to low CR group; 2) within alpha and theta frequency bands, activity will be larger in high than low CR group, mainly during the more difficult task; 3) for global brain activity patterns, we hypothesize that a posterior-to-anterior shift of activity (i.e., a frontalization) will occur in the P300 ERP in low compared to high CR, especially during the more difficult task; 4) another main objective is studying if the posterior-to-anterior shift of activity in older adults with low CR could be associated with a particular frequency band during the tasks and exploring whether it also emerges during the resting state.

## 2. METHODS

### 2.1 Participants

Sixty-seven healthy older adults (30 women) between 61 and 82 years old (M = 70.87, SD = 4.82) took part in the experiment. Participants had normal or corrected to normal visual acuity. The sample was divided into two equal groups by establishing a cut-off for the median value obtained in the CR questionnaire. Thus, participants scoring higher than 15 were included in the high CR group, and participants scoring 15 or less were included in the low CR group. In total, thirty-three older adults were included in the low CR (LCR) group and thirty-four older adults in the high CR (HCR) group. Data from one participant from the HCR group was excluded from time-frequency, resting-state, and frontalization statistical analyses due to technical issues with the processing of the EEG recording. Before taking part in the experiment, all of the participants received information about the experimental procedure and provided their written informed consent.

### 2.2 Procedure

The informed consent was obtained before the beginning of the experimental session. The study was conducted in accordance with the principles of the the Declaration of Helsinki, and ethical permission was given by the local ethics committee. During the first session, the participants took part in a general neuropsychological assessment to make sure that they performed within the normal parameters for age and years of schooling. Namely, participants performed the Mini-mental state examination (Folstein et al., 1975) and the Repeatable Battery for the Assessment of Neuropsychological Status (Randolph et al., 1998). Also, the level of CR was estimated by using a standardized CR questionnaire (Rami et al., 2011, for the English translation of the questionnaire, see Cespón et al., 2023, annex 1). During the second session, participants were in the EEG shielded room. Participants were seated in a comfortable chair while the electrode cap was placed and the two experimental tasks – i.e., the Simon task and spatial Stroop tasks – were performed during the EEG recording. The resting state included EEG acquisition with eyes opened and closed, each one lasting five minutes. The first experimental session (neuropsychological assessment) lasted slightly less than one hour and the second experimental session lasted about 2 hours.

### 2.3 Tasks

As specified before, participants performed two spatial SRC tasks – the more difficult spatial Stroop task and the easier Simon task – while EEG was recorded. Participants were instructed to respond as fast as possible to the color of the stimulus (red or blue) in the Simon task and to the direction pointed by the arrow in the spatial Stroop task by pressing one of the two response buttons arranged horizontally on the keyboard. The tasks are graphically represented in Figure 1.

**Figure 1.**
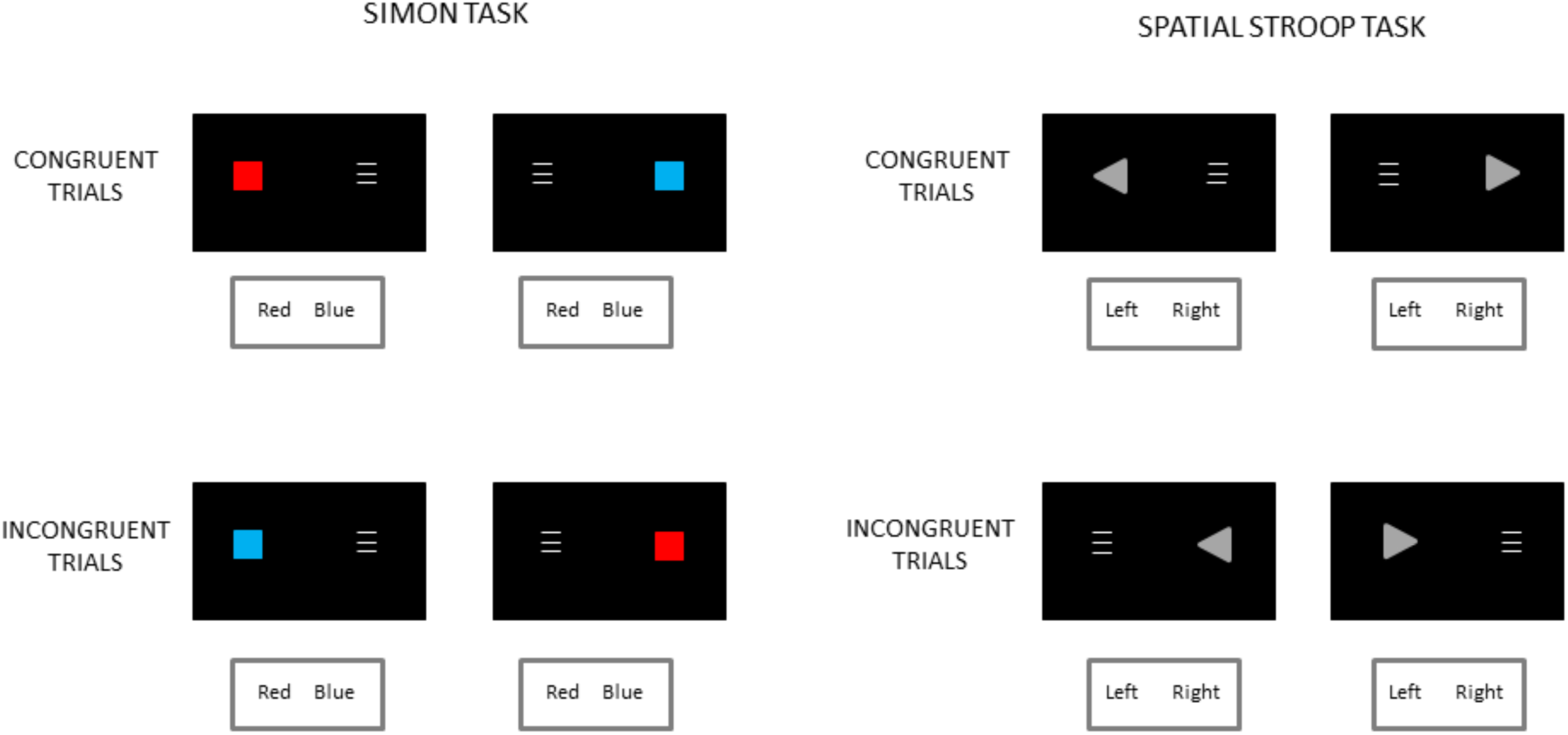
Graphical representation of the Simon and spatial Stroop tasks. *Note.* The top panel represents congruent trials, and the bottom panel represents incongruent trials for both tasks.

During the spatial Stroop and the Simon tasks (Simon & Small, 1969; Cespón et al., 2020; Lu & Proctor, 1995), participants directed their gaze to the center of the screen. A fixation cross appeared in the screen center for 500ms on a black background, followed by an array of stimuli for 100ms. Participants sat 100 cm in front of the screen. The stimuli were presented 5cm far from the central fixation cross so that the entire display was shown within the foveal region (Bargh & Chartrand, 2000). The presence of a contralateral non-target stimulus prevented the influence of asymmetrical N100 activity in central regions (Oostenveld et al., 2001; Valle-Inclán, 1996) and maintained spatial SRC effects (O’Leary & Barber, 1993). Following the presentation of the stimuli, the screen remained blank for a period of 2000±250 ms. The new trial started when the fixation cross appeared in the center of the screen. For each experimental task, 240 trials per condition were presented (i.e., 240 congruent trials, 240 incongruent trials), which resulted in 480 trials. Each task was divided into three blocks of 160 trials per block. At the end of each block, the participant rested for about one minute. Before starting the task, participants completed a practice block of 10 trials.

### 2.4 EEG recording

The continuous EEG was recorded using Easycap (Brain Products GmbH, Germany). The following fifty-four EEG electrodes were placed on the scalp, in accordance with the 10-20 international system: AF7, F7, FT7, AF3, AFz, Fz, F1, F3, F5, FCz, FC1, FC3, FC5, C5, C3, C1, Cz, CP5, CP3, CP1, P7, P5, P3, Pz, CPz, PO7, PO3, O1, Oz, POz, O2, PO8, PO4, P4, P8, P6, CP2, CP4, CP6, C2, C4, C6, FC2, FC4, F4, F2, FC6, FT8, F6, F8, AF4, AF8, T7, T8. Simultaneously to the EEG recording, supraorbital-infraorbital and canthus externus electrodes montages were used to record electro-oculogram (EOG) activity in order to correct artifacts related to ocular movements. Fpz was chosen as the location for the ground electrode. All electrodes were online referenced using the right mastoid. The left mastoid electrode was used offline to re-reference the scalp recordings to the average of the left and the right mastoid. The EEG signal was acquired using a 0.1-500 Hz bandpass filter and digitized at 1000 Hz sampling rate. Impedance was maintained below 5 kΩs.

### 2.5 Data analysis

For analyzing behavioral performance, RTs and response accuracy (Number of Errors – NE) were calculated for each participant. RTs slower than 2000 ms were excluded from the analyses.

The EEG signal was filtered with a 0.1-80 Hz digital bandpass filter and a 50 Hz notch filter. EEG ocular and muscular artifacts were removed from the scalp electrodes by using Independent Component Analysis (ICA) in the BrainVision Analyzer 2 (Brain Products, GmBh, Gilching, Germany). The epochs ranged from −200 to 800 ms, for a total duration of 1000 ms. Epochs that exceeded 100 μV were automatically rejected. All remaining epochs displaying artifacts were also eliminated from further analysis. The baseline was corrected in the interval from −200 ms to 0 ms.

According to previous studies (Fjell & Walhovd, 2001; Guerrero et al., 2022; Speer & Soldan, 2015), ERP latencies and amplitudes for N200 and P300 components were analyzed at representative scalp locations: the Fz (frontal) electrode for N200 (Heidlmayr et al., 2020), and Fz (frontal), Cz (central), and Pz (posterior) electrodes for P300. After inspecting the grand averaged waveforms, we determined latencies with the maximum negative peak between 200 and 350 ms for N200, and the maximum positive peak between 300 and 600 ms for P300. The ERP amplitude was taken ±25 ms around the N200 peak latency (i.e., N200 amplitude was studied as the mean activity in a time window of 50 ms) and ±50 ms around the P300 peak latency (i.e., P300 amplitude was analyzed as the mean activity in a time window of 100 ms).

For the Simon and spatial Stroop tasks, time-frequency analyses were carried out using the Morlet continuous wavelet transform with Gabor normalization after averaging the trials for each subject within the time windows associated with the N200 (200-350 ms) and P300 (300-600 ms) ERP components. We obtained the power (in µV²) within the theta (4.1–7.9 Hz), alpha (8.1–13.9 Hz), and beta (15.1–24.6 Hz) frequency bands within the mentioned time windows at Fz (frontal), Cz (central), and Pz (posterior) electrodes. For resting state, a Fourier transform was used, calculating the spectral power in the main frequency bands (Theta, Alpha, and Beta) in these previously mentioned midline electrodes (i.e., Fz, Cz, and Pz). To obtain indices of frontalization for P300 and spectral activity within each frequency band, we have carried out the following formula: (1+Fz)/(1+Pz). Importantly, we added a constant value of 1 to Fz and to Pz in order to prevent distorted outcomes as a result of using values between 0 and 1 in the division.

### 2.6 Statistical analysis

To analyze RTs and NE, we carried out repeated measures ANOVAs with two within-subject factors: Task (two levels: high difficulty task – i.e., spatial Stroop task- and low difficulty task – i.e., Simon task) and Condition (two levels: congruent, incongruent) and a between-subject factor: Group (two levels: LCR, HCR).

For frontal N200 ERP latencies and amplitudes, repeated measures ANOVAs were carried out with two within-subject factors: Task (two levels: high difficulty task, low difficulty task) and Condition (two levels: congruent, incongruent) and a between-subject factor: Group (two levels: LCR, HCR). For P300 ERP latencies and amplitudes we carried repeated measures ANOVAs with three within-subject factors: Task (two levels: high difficulty task, low difficulty task), Condition (two levels: congruent, incongruent), and Electrode (three levels: Fz, Cz, and Pz), and a between-subject factor: Group (two levels: LCR, HCR).

For alpha, beta, and theta frequency bands, we conducted repeated measures ANOVAs (separately for Fz, Cz, and Pz electrodes) with two within-subject factors: Task (two levels: high difficulty task, low difficulty task), and Congruence (two levels: congruent, incongruent), and a between-subject factor: Group (two levels: LCR, HCR). We used repeated measures ANOVA to analyze the resting state on each studied frequency band (i.e., alpha, theta, and beta) with a within-subject factor: Electrode (Fz, Cz, and Pz), and a between-subject factor: Group (LCR, HCR). Additionally, we utilized independent samples t-tests to explore differences between HCR and LCR groups in frontalization. Moreover, a Pearson correlation analysis was conducted to examine the correlation between frontalization revealed by ERP and spectral activity.

All statistical analyses were carried out using SPSS statistical software version 26 (IBM Corporation, Armonk, NY, USA). The statistical significance was determined by using an alpha level of 0.05. When ANOVAs revealed significant effects, pairwise comparisons were performed using the Bonferroni correction. Partial eta squared (η²p) was used to provide a measure of the effect size.

## 3. RESULTS

### 3.1 Behavioral results

The repeated measures ANOVA (Group × Task × Congruence) for RT (see top part of the Figure 2 and Table 1) revealed a Congruence effect [F (1, 65) = 280.55, p < 0.001, η²p = 0.81], as the RT was slower in incongruent than congruent trials. Moreover, a Task × Congruence interaction effect was significant [F (1, 65) = 37.25, p < 0.001, η²p = 0.36]; specifically, RT was slower in incongruent than congruent trials in the spatial Stroop task (p < 0.001) as well as in the Simon task (p < 0.001). The results showed a Task effect [F (1, 65) = 30.78, p < 0.001, η²p = 0.32]. In detail, RT was slower in the spatial Stroop task compared to the Simon task. The Group factor showed a non-significant trend [F (1, 65) = 3.62, p = 0.06, η²p = 0.05] to slower RT in LCR compared to HCR group.

**Figure 2.**
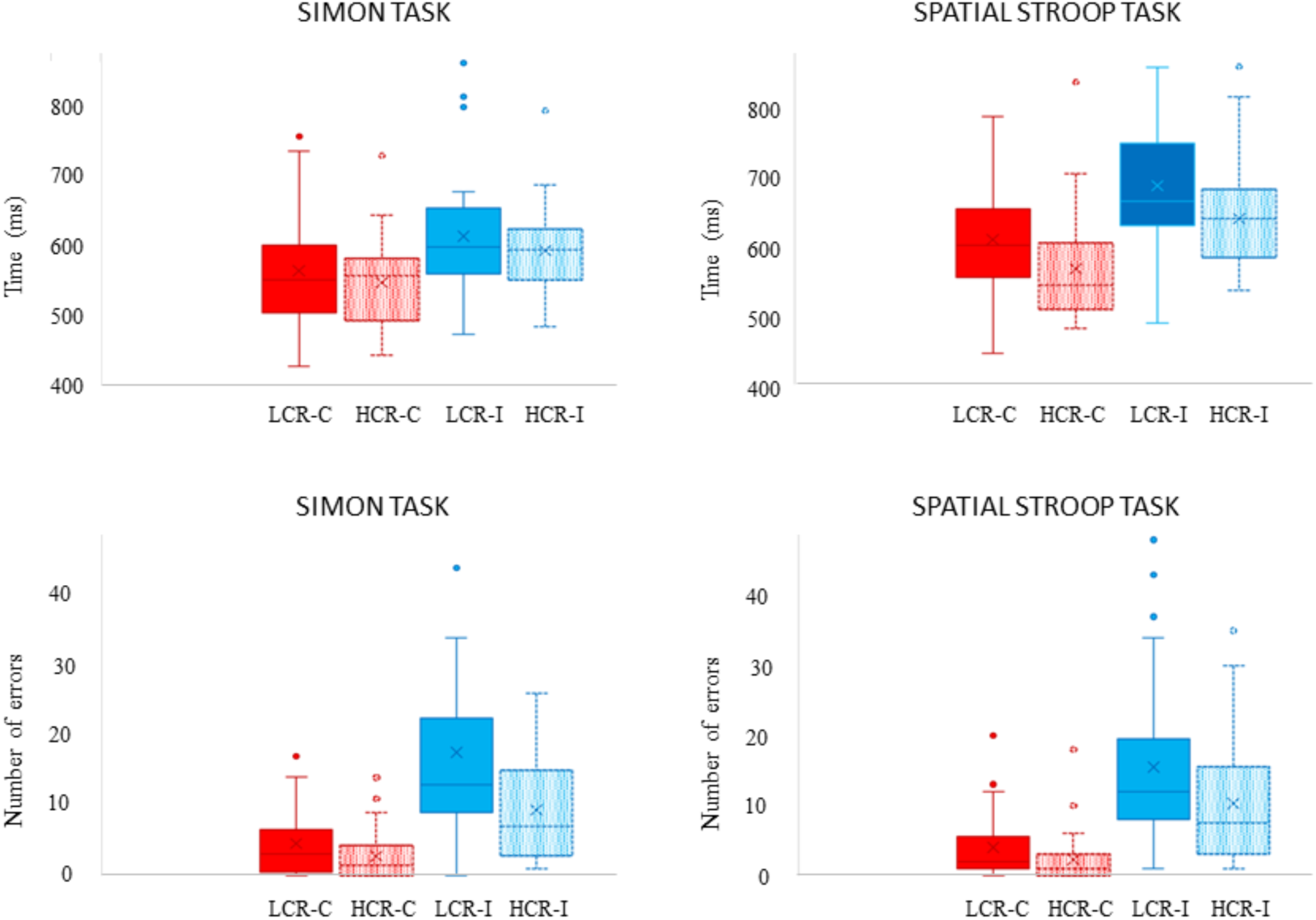
Representation of the obtained behavioral data. *Note.* Mean reaction times and error rates for congruent and incongruent trials in the Simon and Spatial Stroop tasks are compared between high and low cognitive reserve groups. The high cognitive reserve group showed better performance with fewer errors in incongruent conditions compared to the low cognitive reserve group.

**Table 1.** Response Times and Accuracy in Spatial Stroop and Simon Tasks: Comparison Between High and Low Cognitive Reserve Groups.

| Group |  | RT (ms) \| NE (mean) |
| --- | --- | --- |
| LCR |  | 616.17 (11.74) \| 10.38 (1.06) |
| HCR |  | 584.80 (11.57) \| 6.16 (1.05) |
| Task |  |  |
| Simon task |  | 577.57 (8.63) \| 8.56 (0.84) |
| Spatial Stroop Task |  | 623.40 (9.77) \| 7.98 (0.80) |
| Congruence |  |  |
| Congruent |  | 570.02 (7.90) \| 3.37 (0.44) |
| Incongruent |  | 630.95 (8.95) \| 13.17 (1.18) |
| Task x Congruence |  |  |
| Simon task | Congruent | 553.78 (8.50) \| 3.64 (0.50) |
|  | Incongruent | 601.36 (9.04) \| 13.49 (1.33) |
| Spatial Stroop Task | Congruent | 586.26 (9.24) \| 3.11 (0.52) |
|  | Incongruent | 660.54 (10.88) \| 12.86 (1.28) |
| Group x Congruence |  |  |
| LCR | Congruent | 584.59 (11.26) \| 4.24 (0.62) |
|  | Incongruent | 647.75 (12.75) \| 16.53 (1.69) |
| HCR | Congruent | 555.45 (11.09) \| 2.51 (0.61) |
|  | Incongruent | 614.15 (12.56) \| 9.82 (1.66) |

The repeated measures ANOVA (Group × Task × Congruence) for NE (see bottom part of the Figure 2 and Table 1) revealed a Group effect [F (1, 65) = 7.92, p < 0.01, η²p = 0.10], as the NE was higher in LCR compared to HCR group. Congruence also revealed a significant effect [F (1, 65) = 99.41, p < 0.001, η²p = 0.60], as the NE was higher in incongruent than congruent trials. Furthermore, it was observed a significant Group × Congruence interaction effect [F (1, 65) = 6.41, p = 0.01, η²p = 0.09], as the incongruent condition led to higher NE in LCR than HCR (p < 0.01).

### 3.2 ERP results

The repeated measures ANOVA (Group × Task × Congruence) for N200 peak latencies did not show any significant effect. The repeated measures ANOVA (Group × Task × Congruence) for N200 amplitudes revealed an effect of Congruence [F (1, 65) = 11.26, p = 0.001, η²p = 0.14]. In detail, N200 amplitude was larger in incongruent than in congruent trials. Moreover, the ANOVA revealed a significant Task × Group interaction [F (1, 65) = 4.43, p = 0.03, η²p = 0.06], showing that LCR individuals had smaller N200 amplitudes in the spatial Stroop than in the Simon task (p = 0.02). N200 is represented in the Figure 3 and amplitude values are provided in the Table 2.

**Figure 3.**
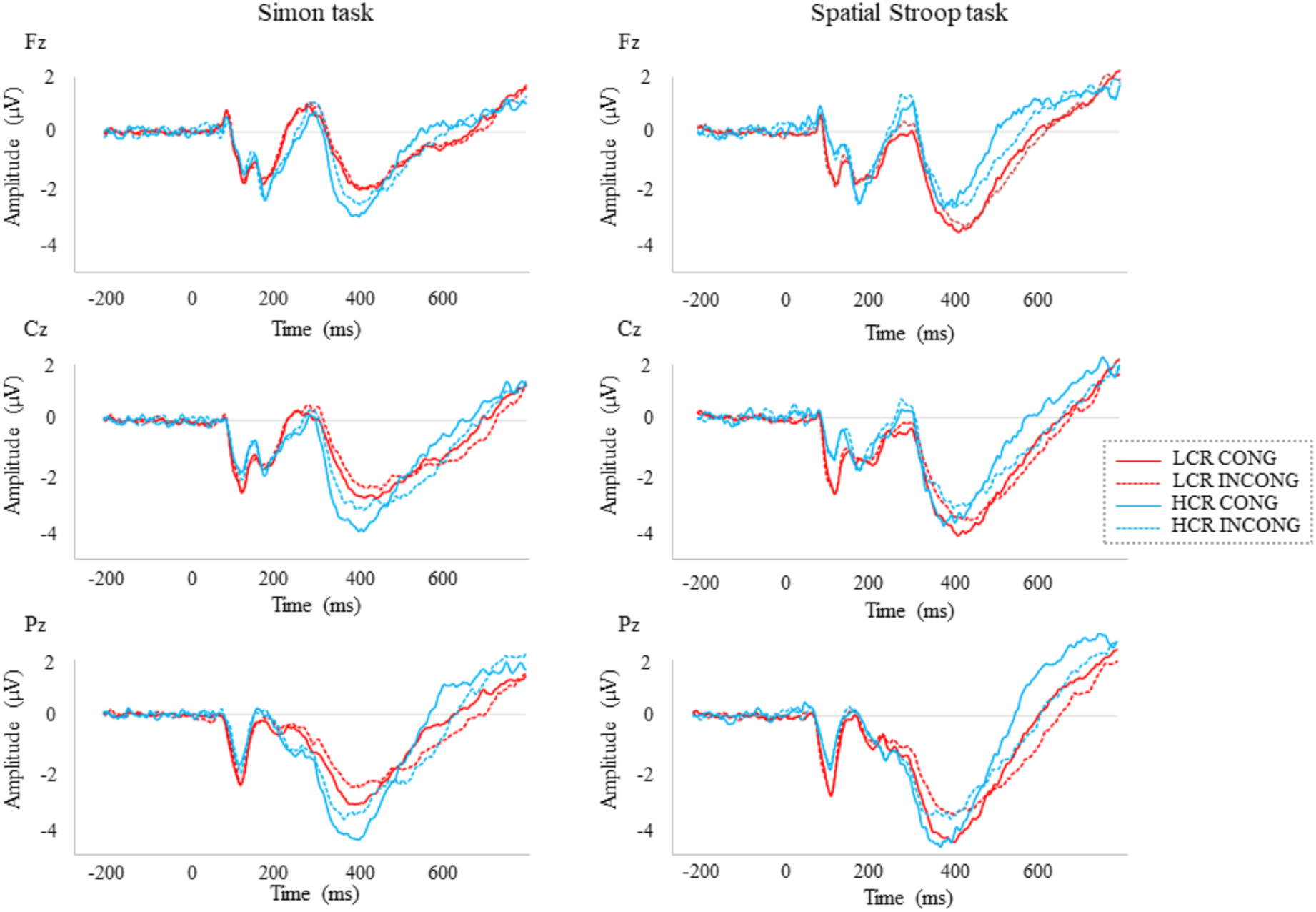
Event-related brain potentials during the executive control tasks. *Note.* Waveforms for congruent and incongruent trials in the Simon and Spatial Stroop tasks, recorded at frontal (Fz), central (Cz), and parietal (Pz) midline electrodes, are compared between high and low cognitive reserve groups. P300 latency occurred earlier in the high cognitive reserve group compared to the low cognitive reserve group.

**Table 2.** N200 Component Characteristics: Mean Amplitudes (μV) for Spatial Stroop and Simon Tasks by Cognitive Reserve Group.

| Group | | Peak Amplitudes for N200 ($\mu V$) |
| --- | --- | --- |
| LCR |  | -1.06 (0.30) |
| HCR |  | -1.31 (0.29) |
| Task |  |  |
| Simon task |  | -1.30 (0.24) |
| Spatial Stroop Task |  | -1.07 (0.23) |
| Congruence |  |  |
| Congruent |  | -1.05 (0.20) |
| Incongruent |  | -1.32 (0.22) |
| Task x Group |  |  |
| Simon task | LCR | -1.39 (0.34) |
|  | HCR | -1.21 (0.33) |
| Spatial Stroop Task | LCR | -0.72 (0.32) |
|  | HCR | -1.42 (0.32) |

The repeated measures ANOVA (Group × Task × Congruence × Electrode) for P300 peak latencies revealed an effect of Group [F (1, 65) = 5.02, p = 0.02, η²p = 0.07], as P300 latency was earlier in HCR than LCR (see Figure 3). The Task × Congruence interaction showed a significant effect [F (1, 65) = 4.19, p = 0.04, η²p = 0.06], as P300 latency was earlier for congruent trials than incongruent trials in the spatial Stroop task (p < 0.01) as well as in congruent trials than incongruent trials in the Simon task (p < 0.001). P300 latency was earlier in congruent than incongruent trials [F (1, 65) = 33.63, p < 0.001, η²p = 0.34].

The repeated measures ANOVA (Group × Task × Congruence × Electrode) for P300 amplitudes revealed a Congruence effect [F (1, 65) = 9.04, p < 0.01, η²p = 0.12] because the P300 amplitude was larger in congruent than incongruent condition. Also, Electrode displayed a significant effect [F (2, 130) = 25.46, p < 0.001, η²p = 0.28], with a smaller P300 amplitude in Fz compared to Cz (p < 0.001) and Pz electrodes (p < 0.001). Task × Group interaction effect was also significant [F (1, 65) = 5.35, p = 0.02, η²p = 0.07], indicating that LCR had larger peak amplitudes in the spatial Stroop than in the Simon task (p < 0.01). The Congruence × Electrode interaction was significant [F (2, 130) = 11.87, p < 0.001, η²p = 0.15]. Specifically, for congruent trials, P300 amplitude was larger in Pz than in Fz (p < 0.001) and Cz electrodes (p = 0.04). For incongruent trials, P300 amplitude was larger in Pz than Fz (p < 0.001) and in Cz than Fz (p < 0.001). The mean values for the P300 peak amplitudes and latencies are represented in Table 3.

**Table 3.** P300 Component Characteristics: Mean Amplitudes (μV) and Latencies (ms) for Spatial Stroop and Simon Tasks by Cognitive Reserve Group.

| Group | | Peak Amplitudes ($\mu\text{V}$) \| Latencies (ms) for P300 |
| --- | --- | --- |
| LCR |  | 3.53 (0.41) \| 441.68 (7.38) |
| HCR |  | 3.60 (0.40) \| 418.46 (7.27) |
| Electrode |  |  |
| Fz |  | 2.90 (0.30) \| 429.07 (6.38) |
| Cz |  | 3.76 (0.32) \| 445.85 (6.38) |
| Pz |  | 4.05 (0.29) \| 415.29 (6.65) |
| Congruence |  |  |
| Congruent |  | 3.70 (0.30) \| 417.98 (5.32) |
| Incongruent |  | 3.44 (0.28) \| 442.16 (5.83) |
| Task x Group |  |  |
| Simon task | LCR | 2.98 (0.42) \| 443.44 (10.30) |
|  | HCR | 3.68 (0.41) \| 422.78 (10.15) |
| Spatial Stroop Task | LCR | 4.08 (0.48) \| 439.92 (7.91) |
|  | HCR | 3.53 (0.48) \| 414.14 (7.79) |
| Task x Congruence |  |  |
| Simon task | Congruent | 3.44 (0.30) \| 417.33 (7.14) |
|  | Incongruent | 3.23 (0.29) \| 448.90 (8.22) |
| Spatial Stroop Task | Congruent | 3.96 (0.37) \| 418.63 (6.02) |
|  | Incongruent | 3.65 (0.32) \| 435.43 (6.45) |
| Congruence x Electrode |  |  |
| Congruent | Fz | 2.93 (0.31) \| 422.52 (7.20) |
|  | Cz | 3.87 (0.33) \| 429.85 (6.30) |
|  | Pz | 4.29 (0.32) \| 401.57 (6.09) |
| Incongruent | Fz | 2.86 (0.29) \| 435.63 (6.54) |
|  | Cz | 3.65 (0.31) \| 461.86 (7.83) |
|  | Pz | 3.80 (0.27) \| 429.00 (8.46) |

### 3.3 ERSP results

For alpha activity, the repeated measures ANOVA (Group × Task × Congruence) at 200-350 ms time window in the Fz electrode showed a main effect of Group [F (1, 64) = 7.35, p < 0.01, η²p = 0.10], indicating higher alpha in HCR than LCR group. Similarly, at 300–600 ms time window in the Fz electrode, alpha activity was higher in HCR compared to LCR group [F (1, 64) = 7.17, p < 0.01, η²p = 0.10]. When examining alpha activity at 300–600 ms time window in the Pz electrode, the analysis showed higher alpha activity in the HCR than in the LCR group [F (1, 64) = 8.38, p = 0.005, η²p = 0.11] (see Figure 4 and Figure 5). The mean values for alpha, beta, and theta are represented in Table 4.

**Figure 4.**
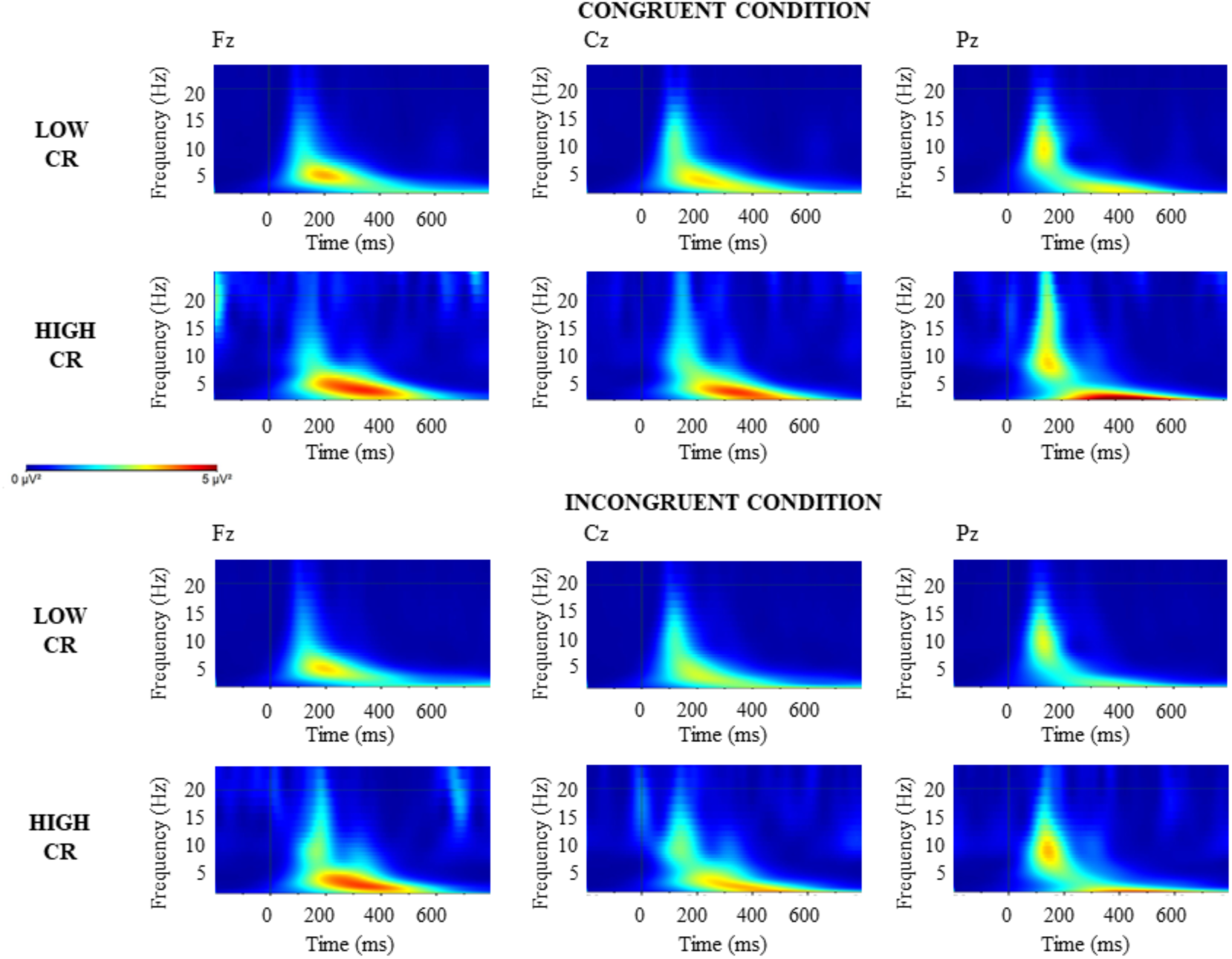
Time-frequency spectral power during the Simon task. *Note.* Time-frequency spectral power during the Simon task. Spectrograms for power changes for congruent and incongruent trials, recorded at frontal (Fz), central (Cz), and parietal (Pz) midline electrodes, are compared between high and low cognitive reserve groups. High cognitive reserve group exhibited greater alpha activity compared to low cognitive reserve group during N200 and P300 time windows in the Simon task.

**Figure 5.**
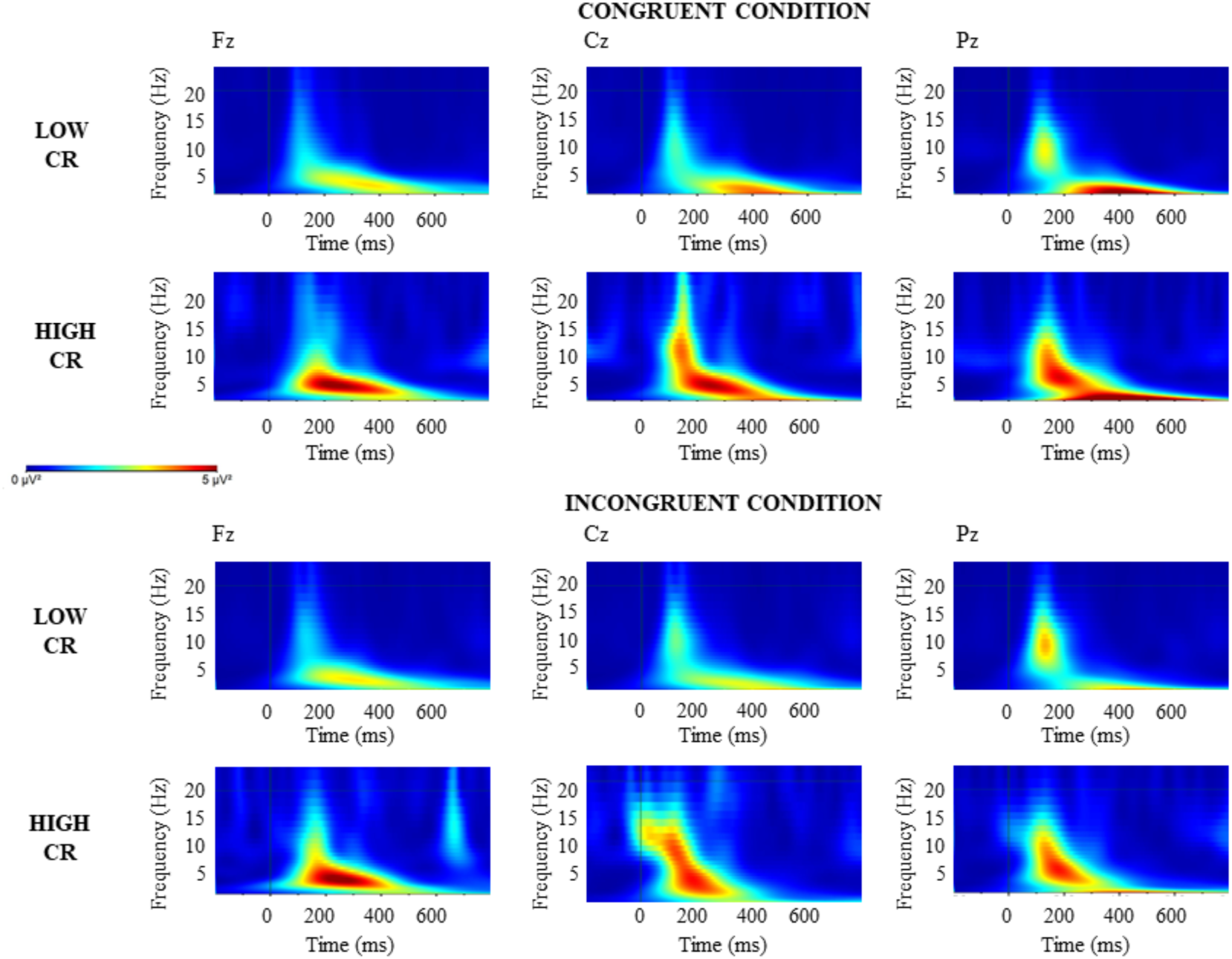
Time-frequency spectral power during the spatial Stroop task. *Note.* Time-frequency spectral power during the spatial Stroop task. Spectrograms for power changes for congruent and incongruent trials, recorded at frontal (Fz), central (Cz), and parietal (Pz) midline electrodes, are compared between high and low cognitive reserve groups. High cognitive reserve group exhibited greater alpha activity compared to low cognitive reserve group during N200 and P300 time windows in the spatial Stroop task.

**Table 4.** Between-Group Differences in Event-Related Spectral Power.

| Spectral power (Hz) in the Fz at a 200–350 ms \| in the Fz at a 300–600 ms \|<br>in the Cz at a 300–600 ms \| in the Pz at a 300–600 ms |  |
| --- | --- |
| Group | Alpha power |
| LCR | 0.59 (0.13) \| 0.20 (0.06) \| 0.19 (0.04) \| 0.18 (0.04) |
| HCR | 1.13 (0.13) \| 0.43 (0.06) \| 0.39 (0.04) \| 0.38 (0.04) |
| Group | Beta power |
| LCR | 2.23 (0.10) \| 0.11 (0.05) \| 0.11 (0.16) \| 0.10 (0.04) |
| HCR | 0.57 (0.10) \| 0.25 (0.05) \| 0.43 (0.16) \| 0.23 (0.04) |
| Group | Theta power |
| LCR | 2.25 (0.48) \| 0.98 (0.17) \| 0.11 (0.16) \| 0.10 (0.04) |
| HCR | 3.31 (0.48) \| 1.44 (0.17) \| 0.43 (0.16) \| 0.23 (0.04) |

For theta activity, the repeated measures ANOVA (Group × Task × Congruence) for theta at a 300–600 ms time window in the Pz electrode found a higher theta in the spatial Stroop task than in the Simon task [F (1, 64) = 4.57, p = 0.03, η²p = 0.06] as well as higher theta for congruent than incongruent trials [F (1, 64) = 6.13, p = 0.01, η²p = 0.08].

For beta activity, the repeated measures ANOVA (Group × Task × Congruence) for beta at 200–350 ms time window in the Fz electrode showed a main effect of the Group [F (1, 64) = 5.02, p = 0.02, η²p = 0.07], with higher beta in the HCR compared to LCR group. The Task × Congruence interaction showed a significant effect [F (1, 64) = 5.04, p = 0.02, η²p = 0.07], indicating higher beta activity in incongruent than in congruent trials in the Simon task (p = 0.03).

### 3.4 Results of spectral power during the resting state

The repeated measures ANOVA (Group × Electrode) for alpha, beta, and theta activity during the resting state (both open eyes and closed eyes conditions) did not reveal any significant main effects or interactions. The spectral analysis at rest is graphically represented in the Figure 6.

**Figure 6.**
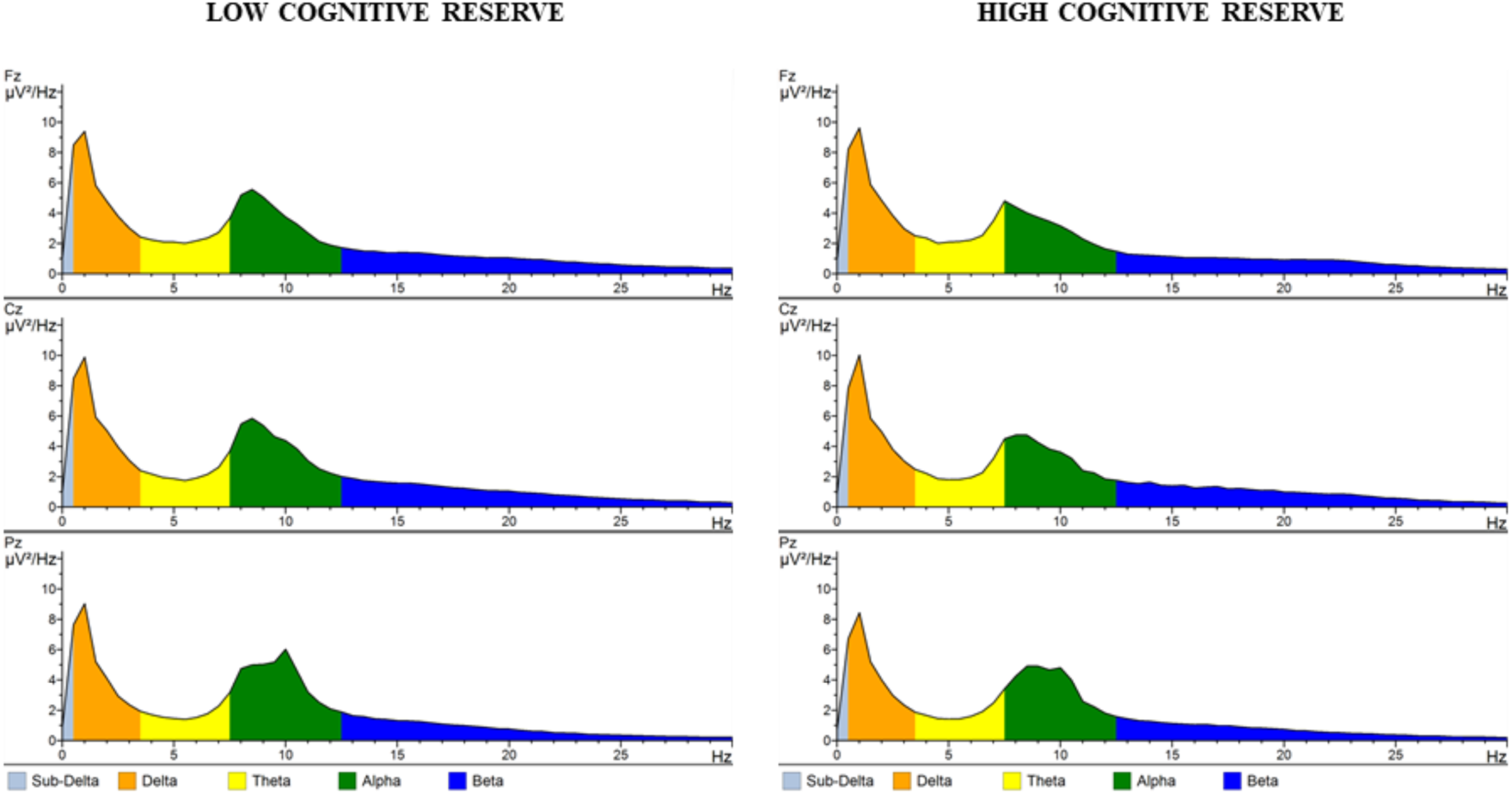
Spectral power during the resting state. *Note.* Spectral power during the resting state. The distribution of spectral power across different frequency bands—delta, theta, alpha, beta, and gamma—during resting-state EEG recordings.

### 3.5 Frontalization indices

For P300, frontalization was higher in LCR compared to HCR group in congruent (0.93 vs. 0.74) [t (64) = 3.29, p < 0.001] and incongruent (1.02 vs. 0.75) [t (64) = 2.88, p < 0.01] conditions of the spatial Stroop task. Differences in the Simon task were not observed. For spectral frequency bands, alpha activity showed higher frontalization in the LCR compared to HCR group in the congruent condition of the Simon task (1.20 vs. 1.02) at 200-350 ms time window [t (64) = 1.69, p = 0.04]. The results from the spectral analysis did not reveal other group-related differences in the calculated frontalization indices.

## 4. DISCUSSION

The results of the present study replicated well-established experimental effects of the SRC tasks, mainly represented by longer RT and higher NE in incongruent than congruent conditions. These effects were related to electrophysiological modulations, which are discussed in subsequent paragraphs. For the main objective of the present study (i.e., investigating differences linked to CR), the results revealed that performance in SRC tasks was better in HCR than in LCR, as displayed by lower NE in HCR compared to LCR group in incongruent conditions. Electrophysiological recordings revealed that P300 latency was earlier in HCR compared to LCR. Furthermore, in the spatial Stroop task, a frontalization of P300 activity was observed in LCR compared to HCR group whereas in the Simon task it was observed a frontalization of alpha activity in LCR compared to HCR group. Also, in LCR group, N200 amplitude was lower in the spatial Stroop task than in the Simon task. Finally, alpha activity was higher in HCR compared to LCR group during the Simon and spatial Stroop tasks. Nevertheless, spectral power differences between HCR and LCR groups in the resting state were not detected in any of the studied frequency bands (i.e., alpha, theta, and beta).

Behavioral results from the current study showed longer RTs in incongruent than congruent conditions as well as increased NE in incongruent compared to congruent conditions for both tasks. This is a well-established behavioral effect of spatial interference, which has been widely demonstrated in spatial SRC tasks (Cespón et al., 2020; Craft & Simon, 1970; Hasbroucq & Guiard, 1992; Juncos-Rabadán et al., 2008; Leuthold & Schröter, 2006; Simon & Small, 1969), and other SRC paradigms (Glaser & Glaser, 1982; MacLeod, 1991). Additionally, we found slower RTs in the spatial Stroop than in the Simon task. These findings are consistent with previous findings that increased task difficulty in the spatial Stroop task leads to participants’ poorer performance (Juncos-Rabadán et al., 2008; Lu & Proctor, 1995; Zhang et al., 1999). An important behavioral finding was that the HCR group had higher accuracy than the LCR group, indicating better cognitive control performance. Importantly, the HCR group showed higher accuracy compared to LCR group in incongruent conditions for both tasks, which is consistent with studies showing increased cognitive control in high compared to low CR older adults (Oosterman et al., 2021; Roldán-Tapia et al., 2012; Yang & Lin et al., 2020).

In the following paragraphs, we briefly discuss the electrophysiological modulations related to the experimental conditions and task difficulty and subsequently we will focus on the main objective of the current research; to know, the electrophysiological differences between LCR and HCR across the three different cognitive states: resting state, performance of the low cognitive control demanding task (i.e., the Simon task) and performance of the high cognitive control demanding task (i.e., the spatial Stroop task).

Electrophysiological modulations related to the experimental conditions showed that P300 latency was later in incongruent than congruent conditions and P300 amplitude was lower in incongruent than congruent conditions of the Simon and spatial Stroop tasks. The results are consistent with previous studies (Leuthold & Sommer, 1999; Leuthold & Schröter, 2006; Melara et al., 2008) and suggest that the time required to update the S-R binding is longer in incongruent than congruent conditions (Adrover-Roig & Barceló, 2010; Cespón et al., 2020). Moreover, lower P300 amplitude was frequently reported in incongruent than congruent conditions of spatial SRC tasks (Cespón et al., 2013; Korsch et al., 2016). This finding is consistent with decreased P300 amplitudes as the task difficulty increases (Polich, 2007) although the P300 latency and amplitude differences between Simon and spatial Stroop task did not reach a significant effect. Another modulation related to experimental conditions of the task was the larger amplitude of the fronto-central N200 in incongruent than congruent conditions, which was usually interpreted as higher cognitive control effort to perform the trials in incongruent compared to congruent conditions (Bartholow et al., 2005; Folstein & Van Petten, 2008; Kopp et al., 1996; Yeung & Cohen, 2006).

For electrophysiological differences related to group, P300 latency was earlier in HCR compared to LCR, which is consistent with the previously formulated hypothesis. This effect was not specific of any task or condition; that is, it affected equally congruent and incongruent conditions as well as Simon and spatial Stroop tasks (i.e., easier, and more difficult cognitive control tasks, respectively). Thus, according to previous studies (Cespón & Carreiras, 2020; Fjell & Walhovd, 2001; Houlihan et al., 1998; Reinvang, 1999), these results may be interpreted as faster neural processing in HCR compared to LCR group. In general, a shorter P300 latency is linked to enhanced cognitive abilities (Polich et al., 1983), which aligns with greater cognitive functioning in HCR than LCR. It is worthy to state that some previous studies observed earlier P300 latencies in high than low CR only in the more demanding tasks (Martinez et al., 2022; Speer & Soldan, 2015) but it is possible that Simon as well as spatial Stroop tasks were both demanding enough to produce the reported P300 differences. N200 latency showed no significant differences between high and low CR groups.

For ERP amplitudes, direct differences between HCR and LCR groups (i.e., a significant effect of the Group factor) were not observed in N200 and P300 amplitudes. However, there was two interesting findings regarding ERP amplitudes that suggest group-related electrophysiological differences in neural processes. Firstly, the N200 amplitudes were reduced during the performance of the spatial Stroop task compared to N200 amplitudes during the Simon task in the LCR but not in the HCR group. Secondly, frontalization of P300 amplitude was observed in LCR compared to HCR in the spatial Stroop task. Next, we discuss the mentioned ERP amplitude modulations.

As previously stated, N200 amplitude was not modulated by the experimental task in the HCR group. In contrast, in the LCR group, N200 amplitude was lower in the spatial Stroop task (i.e., in the more difficult task) than in the Simon task. Such decrease in amplitude may be related to higher neural desynchronization underlying increased effort to perform the task (Gajewski et al., 2018; Willemssen et al., 2011). Considering that fronto-central N200 amplitude is reduced with ageing (Cheng et al., 2019), the decrease of N200 amplitude during the performance of the more demanding cognitive control task in LCR but not in HCR may reflect not only increased effort to accomplish the task but also diminished neural processing efficiency in the LCR compared to HCR group. Interestingly, P300 amplitude modulations were consistent with this interpretation because a posterior-to-anterior shift of activity was observed in the spatial Stroop task for the LCR but not for the HCR group. This neural activity pattern is consistent with higher effort to perform the more demanding task in the LCR compared to HCR group (Cabeza, 2018). Recapping, earlier ERP latencies and a trend to higher ERP amplitudes was observed in the HCR compared to LCR group. In addition, ERPs showed increased frontalization in LCR compared to HCR group during the performance of the more demanding task. These findings fulfill with the view of successful cognitive ageing as related to preserved cognition and maintenance of youthful neural activity patterns (Cespón et al., 2023; Koen & Rugg, 2019; Morcom & Henson, 2018).

The analysis of spectral frequency bands at rest and of spectral time/frequency activity during the performance of the tasks allowed to examine differences related to CR at different brain states (i.e., resting state vs. tasks performance). During the resting state, differences between LCR and HCR were not detected in theta, alpha, or beta frequency bands. On the other hand, during the performance of the Simon and the spatial Stroop tasks, alpha activity was higher in HCR compared to LCR older adults within N200 and P300 time windows. These results allow stating that high levels of CR prevent decrease of alpha activity related to ageing (Barry & De Blasio, 2017) although, in contrast to our hypothesis, effects of CR on theta activity were not observed. In line with the obtained alpha activity results, research exploring the relationship between cognitive performance and alpha activity demonstrated a positive association between higher alpha activity levels and better cognitive performance in older adults (Roca-Stappung et al., 2012; Zangrossi et al., 2021). In short, the current results suggest that functional activity differences between high and low CR emerge more clearly when activating cognitive control networks than during the resting state even if some recent studies (e.g., Varela-López et al., 2022) showed differences between high and low CR during the resting state.

Spectral activity results regarding frontalization were somewhat intricated. Increased frontalization was observed in LCR compared to HCR in alpha activity during the Simon but not during the spatial Stroop task, which contrasts with results observed for the P300 ERP component. A possible explanation might be that, within alpha rhythms, frontalization occurs during the relatively easy tasks whereas there is a loss of frontalization during the more difficult tasks. It is worthy to mention that activity within beta frequency band was higher in HCR than LCR at frontal sites. This has been an unexpected result considering that, in general, beta activity increases with ageing (Barry & De Blasio, 2017).

The main limitation of the current study is that the LCR group included individuals with CR levels ranging from low to middle, attenuating the possible behavioral and neural differences between high and low CR. Ideally, future studies should consider this limitation and recruit older adults with low CR levels to reveal more group-related differences. Another possible direction for future research is to investigate how differences between HCR and LCR groups are modulated by task difficulty by increasing the number of difficulty levels within a given task paradigm (i.e., to design versions of a task with three or more difficulty levels) and/or using other executive control paradigms (e.g., N-back, Go/No-Go tasks). Finally, we did not find any difference between HCR and LCR groups at rest. Our findings strongly suggest that the activation of cognitive control networks is critical for detecting neural correlates of CR. Even so, future studies may focus on neural networks by carrying out functional connectivity analyses comparing HCR and LCR groups at rest and during the task’s performance. For this purpose, such studies should optimize the experimental procedures by taking into account the methodological caveats existing in EEG functional connectivity analysis (Bastos & Schoffelen, 2016).

## 5. CONCLUSIONS

The main objective of the present study was to identify behavioral and neural differences between LCR and HCR groups as well as investigating to what extent such differences may change during the resting state, the performance of a low demanding cognitive control task (i.e., Simon task) and the performance of a high demanding cognitive control task (i.e., spatial Stroop task). The results of the present study showed better cognitive control in HCR compared to LCR group, as revealed by higher accuracy in incongruent conditions of both tasks. Several electrophysiological differences between HCR and LCR groups were observed during the performance of the cognitive control tasks, but electrophysiological differences were not detected during the resting state. Importantly, electrophysiological differences were more pronounced between HCR and LCR groups during the more demanding task (i.e., the spatial Stroop task). These results suggest that differences between high and low CR increase when cognitive control networks need to be substantially activated to cope with task demands. Also, electrophysiological activity during the performance of the cognitive control tasks show that older adults with high CR display neural activity patterns typically observed in younger adults, which is supported by the following findings: 1) earlier P300 latency in HCR than LCR; 2) decreased N200 amplitude during the performance of the spatial Stroop task compared to Simon task in LCR but not in HCR; 3) higher alpha evoked-activity in HCR compared to LCR; 4) frontalization of several electrophysiological correlates (P300 and alpha) in LCR compared to HCR. Overall, the results of the present study show that, in cognitively healthy older adults, high levels of CR relate to maintenance of neural activity patterns observed in younger adults rather than deployment of neural compensatory mechanisms.

## DECLARATIONS

### Funding

This project has received funding from the Spanish Ministry of Science (JC: PID2019-105538RA-I00), Ramón y Cajal program (JC: No. RYC2022-035443-I), from the Basque Government through the BERC 2022-2025 program, and from the Agencia Estatal de Investigación through BCBL’s Severo Ochoa excellence award CEX2020-001010-S. The funding sources were not involved in any activity contributing to this publication.

### Conflicts of interest

The authors report that they have no relevant financial or non-financial interests to declare.

### Ethics approval

The study was conducted in accordance with the principles of the Declaration of Helsinki. Approval was obtained from the Ethics Committee of the Basque Center on Cognition, Brain and Language.

### Consent to participate

Written informed consent was obtained from all individual participants included in the study.

### Consent for publication

Not applicable.

### Availability of data and materials

Data are available upon request from the authors. None of the experiments were preregistered.

### Code availability

Not applicable.

### Authors’ contributions

AZ: processed and analyzed the data, results interpretation, wrote the manuscript. MCP: guided data analysis, critical revision of the manuscript. MC: conceptualization of the experimental design, critical revision of the manuscript. JC: conceptualization of the experimental design, guided data analysis, results interpretation, critical revision of the manuscript.

## Notes

### Competing Interest Statement

The authors have declared no competing interest.

